# Patient-specific deep offline artificial pancreas for blood glucose regulation in type 1 diabetes

**DOI:** 10.1101/2022.10.21.513303

**Authors:** Yixiang Deng, Kevin Arao, Christos S. Mantzoros, George Em Karniadakis

## Abstract

Due to insufficient insulin secretion, patients with type 1 diabetes mellitus (T1DM) are prone to blood glucose fluctuations ranging from hypoglycemia to hyperglycemia. While dangerous hypoglycemia may lead to coma immediately, chronic hyperglycemia increases patients’ risks for cardiorenal and vascular diseases in the long run. In principle, an artificial pancreas – a closed-loop insulin delivery system requiring patients manually input insulin dosage according to the upcoming meals – could supply exogenous insulin to control the glucose levels and hence reduce the risks from hyperglycemia. However, insulin overdosing in some type 1 diabetic patients, who are physically active, can lead to unexpected hypoglycemia beyond the control of common artificial pancreas. Therefore, it is important to take into account the glucose decrease due to physical exercise when designing the next-generation artificial pancreas. In this work, we develop a deep reinforcement learning algorithm using a T1DM dataset, containing data from wearable devices, to automate insulin dosing for patients with T1DM. In particular, we build patient-specific computational models using systems biology informed neural networks (SBINN), to mimic the glucose-insulin dynamics for a few patients from the dataset, by simultaneously considering patient-specific carbohydrate intake and physical exercise intensity.

## 1 INTRODUCTION

Diabetes mellitus is a growing epidemic and its prevalence has been increasing in nearly all countries (1). Global estimates show that the prevalence of adults aged 20-79 years is 8.8% in 2015, and it is predicted to rise to 10.8% in 2040 (2). In the US, according to the Centers for Disease Control and Prevention (CDC), a total of 34.2 million people have diabetes or 10.5% of the US population in 2018 (3). The most common forms of diabetes are type 1 and type 2 diabetes mellitus. In 2016, US data showed type 1 and type 2 diabetes accounted for approximately 6% and 91% of all cases of diagnosed diabetes, respectively (4). Type 1 diabetes (T1DM) is due to autoimmune B-cell destruction, which usually leads to absolute insulin deficiency. On the other hand, type 2 diabetes (T2DM) is due to progressive loss of adequate B-cell insulin secretion with associated insulin resistance (5, 6). Glucose is integral in energy consumption as it serves as a primary metabolic fuel. Under normal physiology, in a fasting state, there is a basal insulin secretion to help match hepatic gluconeogenesis to maintain a blood glucose target between 70 and 130 mg/dl. After a meal, there is a rise in blood glucose levels resulting in concomitant increase secretion of insulin from pancreas (7). The major effects of insulin on glucose metabolism are the following: (a) increases glucose transport across the cell membrane in adipose tissue and muscle, (b) increases glycolysis in muscle and adipose tissue, (c) stimulates glycogenesis and inhibits glycogenolysis in muscle, and liver, and (d) inhibits gluconeogenesis in the liver (8). A few hours after the meal, as the blood glucose concentration falls, glucagon is secreted to release glucose back to the blood which decreases glucose fluctuations. The main goal of treatment, especially for T1DM patients, is to mimic these physiologic insulin secretion by providing appropriate basal and prandial insulin doses. Patients with T1DM are frequently suffering from complications associated with unstable glucose levels, when the blood glucose (BG) regulation is not well controlled. According to the Diabetes Control and Complications Trial, better BG control leads to lower HbA1c levels, which finally leads to better outcomes in terms of both microvascular and macrovascular complications (9). Hence, achieving stable glucose levels has been a well established goal in diabetes management.

Unlike a fingerstick test providing one-time reading of blood glucose from a blood sample (10), wearable minimally-invasive continuous glucose monitoring (CGM) sensors can provide real-time blood glucose concentration measurements for days, by measuring from interstitial fluid (11). When integrated with CGM, another implantable or wearable device, namely “insulin pump”, provides continuous insulin administrations through subcutaneous infusion instead of needle injection, greatly improve the quality of life for patients with diabetes (12, 13, 14). Combining a glucose sensor, an insulin infusion device, and a control algorithm, artificial pancreas (AP) is a closed-loop system designed for patients with T1DM to improve their BG regulation, and consequently decrease the risk of diabetic complications (15, 16, 17, 18). While a few control algorithms in the AP design (19, 20), show sufficient performance at the general population level, very few of them address the inter- and intra-patient variability. More importantly, although patients benefit from regular physical activity in terms of the diabetes management (21, 22, 23), the exercise-induced glucose uptakes also leads to negative effect of the pre-programed insulin usage, leading to increased risk of hypoglycemia during physical exercises (24, 25). Unfortunately, most existing AP algorithms consider only limited external factors like meal intakes which increase plasma glucose levels and insulin infusion which lower the glucose levels (26, 27, 28, 29, 30), and some also require meal announcement (31).

Therefore, the next-generation AP in precision medicine requires a holistic design considering features such as the patient-specificity, physical exercises, and less human intervention. Fortunately, rapid growth of artificial intelligence (32, 33), especially deep neural networks based data-driven machine learning algorithms, provides us with opportunities to characterize the complex dynamics in patients’ metabolic environment, and further enhance the development of control algorithms like reinforcement learning. However, regular reinforcement learning algorithms demands large training datasets in the learning process and constant interaction between the learned agent and environment, i.e., patient’s body, in healthcare related tasks. Despite clinical trials providing abundant medical data from the wearable or implantable devices on patients (34, 35), the specific requirement of agent-environment interaction in reinforcement learning undoubtedly is a huge barrier considering the safety of patients during data collection. Fortunately, utilizing only previously collected offline data without additional online interaction with the environment, offline reinforcement learning algorithms may provide a good opportunity to mitigate the potentially high expense and danger of data collection in healthcare related problems (36).

Motivated by the features highlighted for next-generation AP design, we propose a novel framework to design a patient-specific artificial pancreas integrating wearable device data, systems biology informed neural networks and offline reinforcement learning algorithms (Fig. 1). By simultaneously considering patient-specificity, meal intakes, insulin infusion, and most importantly physical exercises, we first build a patient-specific simulator (digital twin) using a system of ordinary differential equations (ODE) developed by Roy and Parker (37), systems biology informed neural networks (SBINN) (38) and wearable sensor data from the OhioT1DM dataset (39, 40). After obtaining the dynamics of hidden states and hidden parameters for the simulator, we implement offline reinforcement learning algorithms on the patient-specific simulator to improve the plasma glucose dynamics with a safer and more efficient insulin dosage planning. In particular, our offline agents learn the insulin dosage with only a short sequence of past glucose levels without any meal or exercise announcements, which significantly further the step towards an authentic closed-loop system for artificial pancreas design. Our results suggest that the proposed framework provides in-depth diagnosis on patients’ diabetes management and leads to an insightful prognosis on medical improvements at a patient-specific level. We are confident that this framework could benefit the design of in silico clinical trials, where some patient-specific data and clinically verified physiological (parametrized) models are available.

**Figure 1:**
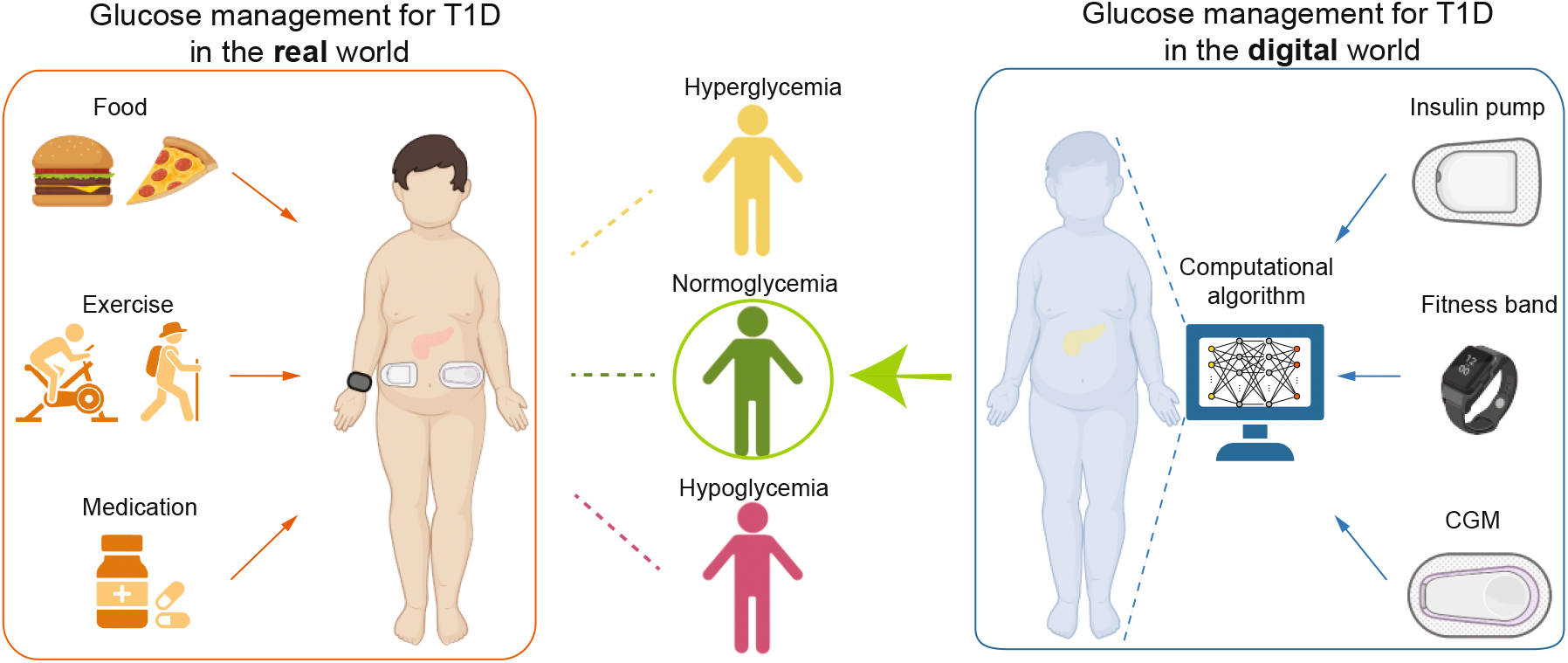
Motivation of digitize blood glucose management in type 1 diabetes. With interaction to the real world, T1D patients experience fluctuating blood glucose from hypoglycemia (BG levels lower than 70 mg/dl, 80 mg/dl if measured by CGM), normoglycemia (BG levels between 80 and 180 mg/dl), and hyperglycemia (BG levels greater than 180 mg/dl). In the digital world, a digital twin mimicking the glucose-insulin dynamics can be created and optimized using computational algorithm with real-world inputs, i.e., insulin pumps provided exogenous insulin along with carbohydrate intakes, fitness bands provided the exercise intensity, and CGM provided real-time glucose levels. BG, blood glucose. CGM, continuous glucose monitoring. Part of the image is created using Biorender.

## 2 METHODS

### 2.1 Framework of the study

In this work, we developed a computational framework to design a patient-specific automated insulin delivery system for six patients with type 1 diabetes using patient-specific data from the OhioT1DM dataset. The OhioT1DM dataset contains eight-week continuous glucose monitoring, insulin, physiological sensor, and self-reported life-event data for 12 patients with type 1 diabetes, among which 6 patients participated in the 2018 cohort and the other 6 patients in the 2020 cohort (40). A workflow of this framework is shown in Fig. 2A. We first implemented systems biology informed neural networks (SBINN) on the OhioT1DM dataset for parameter inference of the patient-specific Roy-Parker model (37), based on historical records of total exogenous insulin (bolus insulin and basal insulin), carbohydrate intakes, heart rate, and CGM measured glucose level. We then trained a deep offline reinforcement learning neural network to build a patient-specific automated insulin delivery system for two representative patients in the OhioT1DM dataset. The final optimized agent, represented by a deep neural networks, can serve as the patient-specific artificial pancreas, leading to a better insulin dosage scheme for the patient.

**Figure 2:**
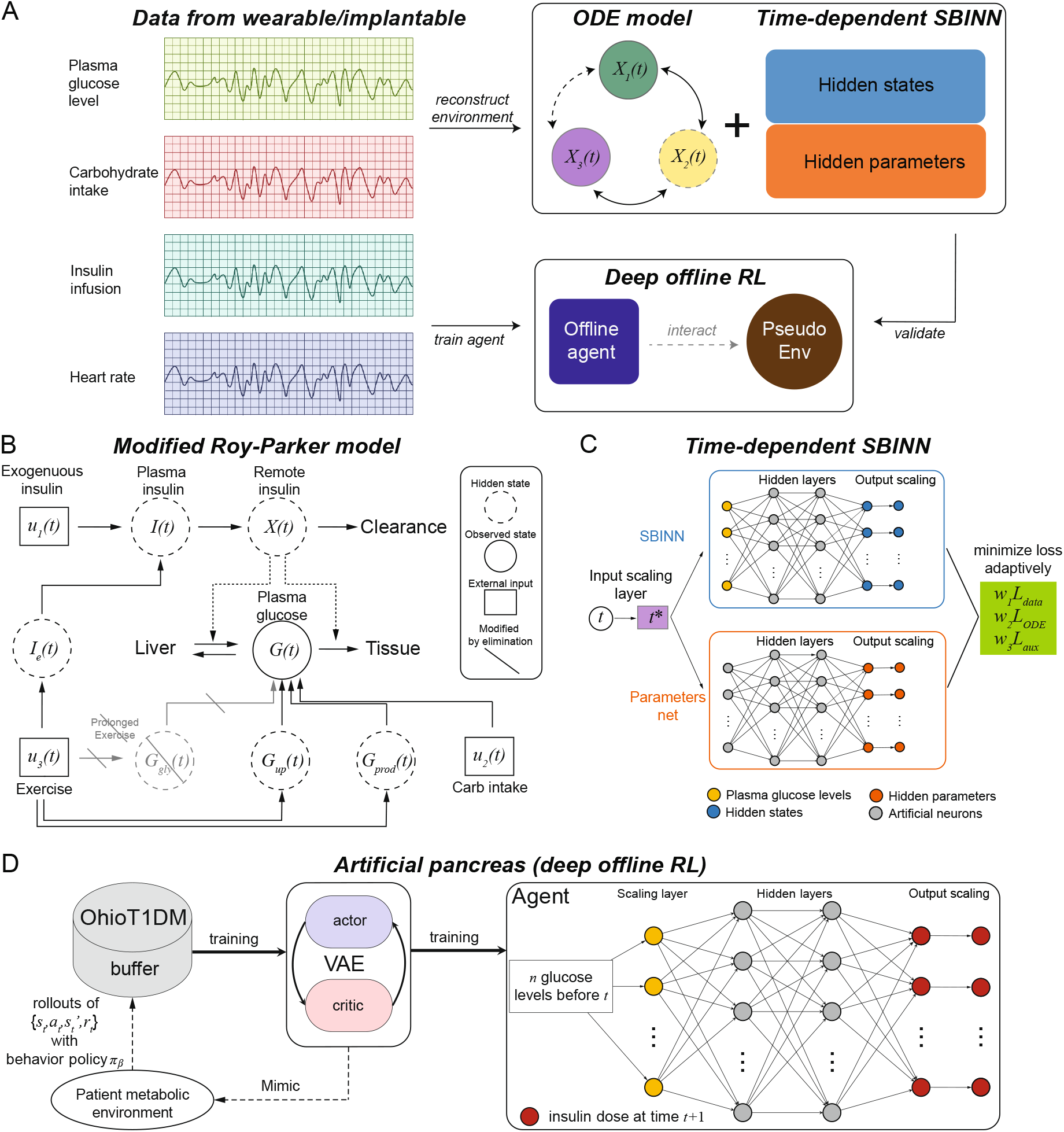
Overview of patient-specific deep offline reinforcement learning for blood glucose regulation. (**A**) Schematic of the framework proposed. Data collected from wearable or implantable devices, such as plasma glucose levels, carbohydrate intakes, insulin infusion and heart rates, were used in two ways. Firstly, the data is applied to reconstruct patient’s metabolic environment, represented by a ODE model and systems biology informed neural networks (SBINN). Meanwhile, the data is also used to train an offline agent composed of neural networks with reinforcement learning algorithms. The reconstructed environment is used to validate the trained offline agent. Part of the image is created using Biorender. (**B**) The compartment diagram of an ODE model, namely modified Roy-Parker model, captures the intrinsic relationships among the observed state *G* and other hidden state variables, considering external inputs like carbonhydrate intakes and physical exercises intensity. (**C**) Schematic of the time-dependent systems biology informed neural networks (*SBINN*) with self-adaptive weights proposed to learning the dynamics of hidden states and the hidden model parameters. (**D**) Schematic of artificial pancreas design using deep offline reinforcement learning, specifically, batch-constrained Q learning (BCQ) algorithm. A buffer derived from the OhioT1DM dataset was used to train a variational autoencorder (VAE), which partially mimics the patient’s metabolic environment in an offline fashion. An agent represented by a deep neural networks was then trained using merely *n* plasma glucose levels before time *t* (*t* included) to provide the insulin dosage at time *t* + 1. By iteratively marching forward, a better insulin dosing plan is generated.

### 2.2 OhioT1DM dataset

#### Dataset overview

All patients in the OhioT1DM were on insulin pump therapy with continuous glucose monitoring (CGM), throughout the 8-week data collection period. During these 8 weeks of monitoring, the patients also reported life-event data via a custom smart-phone app and provided physiological data from a fitness band. Based on the form of the ODE model, i.e., the Roy-Parker model, studied in this work, we selected from the OhioT1DM dataset the following historical measurements: 1) the CGM blood glucose level, 2) insulin doses, both bolus insulin and basal insulin, 3) self-reported meal times and the amounts of carbohydrate intakes, and 4) the heart rate. We note that only the data of those 6 patients in the 2018 cohort is used in our analysis, due to the lack of heart rate monitoring in the 2020 cohort.

#### Data preprocessing

The exogenous insulin is calculated as the sum of basal insulin and bolus insulin at each moment. According to the user manual of the insulin pumps, i.e., Medtronic 530G and 630G, while the basal insulin is given at a rate and is provided explicitly, bolus insulin is a one-time dose and can be released into the blood stream using different mODE, i.e., “normal”, “normal dual”, “square” and “square dual”. Given limited information for the exact releasing process of these different mODE in the OhioT1DM dataset, we assume the conversion formula based on literature (41) and the user guides of the corresponding commercial insulin pumps (42, 43). For “normal” type, the single *x* mU dose bolus insulin is converted to a constant pseudo basal insulin at a rate of *x*/10 mU lasting for 10 minutes. For “square” type, the single *x* mU dose bolus insulin is converted to a constant pseudo basal insulin at a rate of *x/t* mU lasting for *t* minutes, where *t* denotes the time elapse between two adjacent boluses. For “normal dual” and “square dual”, the single dose is divided evenly into two identical half doses. These two half doses are released sequentially by “normal” and “square” type in “normal dual”, while by “square” and “normal” type in “square dual”, respectively. Additionally, we also consider “temp basal” insulin, which overrides the basal insulin set previously. The glucose consumption rate *u*_2_ due to meal intakes is computed by the amount of meal carbohydrate using an exponential decay function (38) as follows,

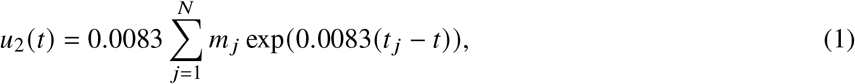

where *m_j_* gram of carbohydrate intake is recorded at *t _j_*, *N* is the total number of meals. The percentage of 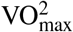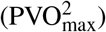 denoting a percentage of the maximum rate of oxygen consumption due to exercise is approximated by the heart rate (HR) as follows (44),

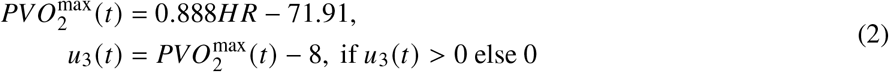

where 8 denotes the average 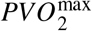 for a person at the basal state (37). After converting all data into the same resolution, we further smoothed the data by using a rolling window and generate a formatted dataset, where the sampling interval is 1 hour between neighbouring time points. Fig. S2 shows the processed insulin infusion *u*_1_ (*t*), carbohydrate intake *u*_2_ (*t*) and exercise intensity *u*_3_ (*t*) for 6 patients in the OhioT1DM dataset.

### 2.3 Systems biology informed neural networks (SBINN) with the Roy-Parker model

#### Roy-Parker model

With the aim of developing a robust closed-loop insulin delivery system under changing physiological conditions, Roy and Parker developed a model that can predict blood glucose levels at rest and during physical exercises (37). Since the patients participated in OhioT1DM only performed sporadic and light physical activities, we modified t he O DE s ystem i n Roy a nd Parker (37) by o mitting t he *G_gly_* (*t*) term representing the decline of the glycegenolysis rate during prolonged exercise due to the depletion of liver glycogen stores (Fig. 2B). The resulting ODE system for the 6 state variables [*I, X, G, G_prod_, G_up_, I_e_*] is shown in Eqs. (3)–(8),

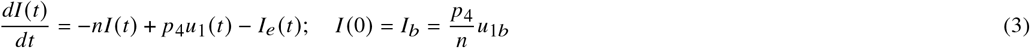

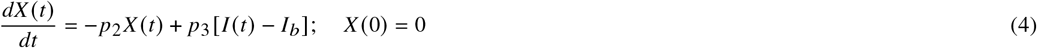

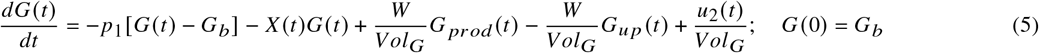

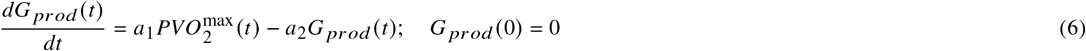

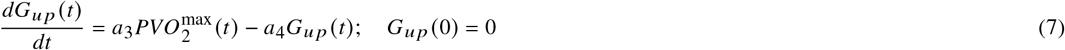

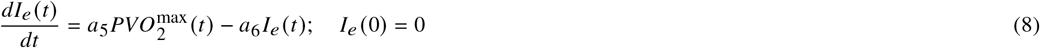

It captures the exercise-induced dynamics of plasma insulin concentration *I* (*t*), remote insulin concentration *X*(*t*), the plasma glucose level *G*(*t*), exercise-induced hepatic glucose production *G_prod_* (*t*), exercise-induced glucose uptake *G_up_* (*t*), exercise-induced insulin removal from the circulatory system *I_e_* (*t*), exogenous infusion *u*_1_ (*t*), and external glucose uptake *u*_2_ (*t*). The instant parameters *I_b_* and *G_b_* represent the basal plasma insulin and glucose concentrations, respectively. Given the multi-scale nature of the glucose levels in a month-long observation, we found that our algorithm learns better the glucose dynamics when we allow the parameters in the ODE to vary over time. Table S1 shows the nomenclature, physiological meaning and reference values of patient-specific parameters to be inferred in the ODE. The ranges of the parameters were set to [0.2·*x_ref_*, 1.8·*x_ref_*], where *x_ref_* represents the corresponding reference value of the variable in Table S1.

#### Systems biology informed neural networks (SBINN)

Yazdani et al. developed a novel systems-biology-informed deep learning framework, namely systems biology informed neural networks (SBINN), which successfully incorporates the system of ordinary differential equations (ODE) into the neural networks (38). Inspired by the physics-informed neural networks, SBINN, shown in Fig. 2C, is sequentially composed of an input-scaling layer to allow input normalization for the robust performance of the neural networks, a feature layer marking different patterns of state variables in the ODE system and the output-scaling layer to convert normalized state variables back to physical units. By effectively adding constraints derived from the ODE system to the optimization procedure, SBINN is able to simultaneously infer the dynamics of unobserved species, external forcing, and the unknown model parameters.

Given the measurements of *y*_1_, *y*_2_,…, *y_M_* at times 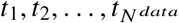, SBINN enforces the network to satisfy the ODE system at the time point 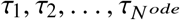. The total loss is defined as a function of both the parameters of the neural networks, denoted by ***θ*** and parameters of the ODE system, denoted by ***p***.

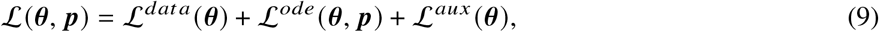

where 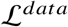 is associated with the *M* sets of observations of the state variables ***y*** in the ODE system; 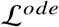 enforces the structure imposed by the system of ODE; 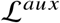 is introduced as an additional source of information for the system identification. We recommend Ref. (38) that provides further details of the loss functions. The final step of SBINN is to infer the neural network parameters ***θ*** as well as the unknown ODE parameters ***p*** simultaneously by minimizing the aforementioned loss function via gradient-based optimizers.

In this work, the known observed state variable *y* is the CGM measured glucose record in the OhioT1DM dataset, i.e., *G*(*t*), which is used for minimizing the data loss, 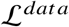. We used 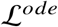 to minimize the residual terms in the ODE, shown in Eqs. (3)–(8). Following Yazdani et al. (38), we imposed the initial condition as the auxiliary loss 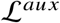. To improve the training of SBINN and speed up the convergence, we implemented self-adaptive weights over each iteration on the weights of each loss terms (45). The self-adaptive weighted loss 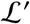 is as follows,

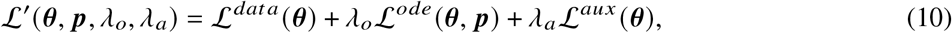

where *λ_o_* and *λ_a_* are trainable, non-negative self-adaptation weights associated to the ODE loss term and auxiliary loss term, respectively. Hence, the objective of the training of neural networks is updated to

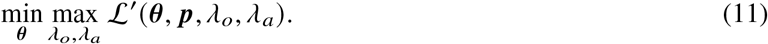

The update rules for the self-adaptive weights are given by,

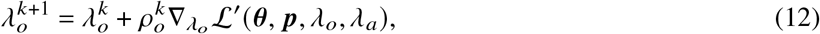

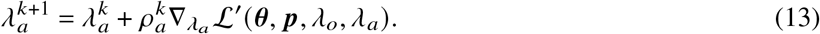

where 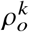 and 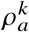 denotes the learning rates for the corresponding weights, which was set to be 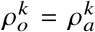 in this work.

### 2.4 Offline reinforcement learning

Unlike supervised learning where a direct input and output mapping is given explicitly, reinforcement learning algorithms define the control problem as a Markov Decision process, and train an agent, which collects a step-by-step reward through interactions with the environment of interest. The goal of the agent is to respond to the environment changes such that the total reward of the series of responses, namely actions, is maximized by the end of training. While (online) reinforcement learning requires a demanding setting where the agent is trained through trial-and-error interactions with a dynamic environment, offline reinforcement learning algorithms utilize previously collected data, with no extra interaction with the environment (36, 46). However, the benefits of offline reinforcement learning also come with some disadvantages, for example the distributional shift, i.e., while the function approximations might be trained under one distribution, it will be evaluated on a different distribution. Efforts to address this challenge can be generally grouped into two categories: (1) policy constraint methods, which constrain the learned policy to lie close to the behavior policy, which was used to collect the offline dataset; and (2) uncertainty based methods, which attempt to estimate the epistemic uncertainty of value functions, and then utilize this uncertainty to detect distributional shift. A comprehensive review of offline reinforcement learning algorithms could be found in (36). Despite of some disadvantages, offline reinforcement learning is a favorable choice for tasks demanding a lot of time and cost when interacting with the environment, for example, healthcare related tasks. As shown in Fig. 2D, in offline reinforcement learning, a buffer dataset is collected by some behavior policy *π_β_*, before the training starts. Afterwards, the agent represented by a deep neural networks is trained with an offline reinforcement learning algorithm, specifically, batch-constrained Q learning (47). Finally, a policy outperforming the behavioral policy *π_β_* is deployed in the patient-specific artificial pancreas. In this work, we generated a buffer by sampling from the OhioT1DM dataset and generate the states, i.e., glucose levels, carbon intakes and physical exercises, along with the action denoted by total exogenous insulin at a specific time point, i.e., a sum of the bolus insulin and basal insulin, and the corresponding returns depending on the resulting glucose levels.

#### Batch-constrained Q learning (BCQ)

Herein, we implemented the batch constrained Q-learning (BCQ) algorithm, which is one of the most popular offline reinforcement learning algorithms (47). By restricting the action space in order to force the agent towards behaving close to a subset of the given data, BCQ is able to learn successfully without interacting with the environment by considering extrapolation error. The details of the BCQ algorithm are shown in Algorithm 1. Following Fujimoto et al. (47), we implemented a fully-connected feed forward neural networks for the Q-networks and the variational autoencoders (VAE). The hyper-parameters used in this work can be found in Table S2.

## 3 RESULTS

To build a surrogate environment for our agent to interact with, we first performed patient-specific parameter inference using SBINN with a system of ODE developed by Roy and Parker (37). The primary step of inference for a system of ODE is to examine its identifiability. After obtaining the patient-specific parameters of the ODE, which is essential to reconstruct the dynamics of the state variables, we developed a patient-specific offline reinforcement learning algorithm to learn an optimal planning for the external insulin infusion for two representative patients, which helped them decrease the risk of hypoglycemia and hyperglycemia, respectively.

### ODE identifiability of the Roy-Parker model

We first perform structural identification on the ODE for the set of parameters

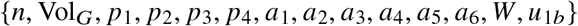

appeared in Eqs. (3)–(8). Although most of the ODE parameters are not readily available in the OhioT1DM dataset, there are several parameters that are practically available for patients. For example, the patient’s body weight (*W*) can be measured practically but is not available in OhioT1DM. While *u*_1*b*_, the exogenous insulin infusion rate to maintain basal plasma insulin, is not available in the OhioT1DM, but it could be inferred from the mode of the exogenous insulin profile of the specific patient. We considered the identifiability of the afore-mentioned parameters under different scenarios depending on whether *W* and *u*_1*b*_ are known for the given patient. Table 1 suggests that when *W* and *u*_1*b*_ are both known, other parameters are either globally identifiable or locally identifiable. We adopted this scenario in the following analysis by assuming *W* = 60 *kg* and patient-specific *u*_1*b*_ to construct the patient-specific model by inferring a subset of the aforementioned parameters {*n*(*t*), Vol_*G*_(*t*), *p*_1_(*t*), *p*_2_(*t*), *p*_3_(*t*), *p*_4_(*t*), *a*_1_(*t*), *a*_2_(*t*), *a*_3_(*t*), *a*_4_(*t*), *a*_5_(*t*), *a*_6_(*t*)} for one-month period.

**Table 1:** Parameter identifiability in the Roy-Parker model under four different scenarios. *W* denotes the bodyweight of the patient, *I_b_* denotes the basal plasma insulin level. Details of other parameters can be found in (37).

| Parameter | Identifiability |  |  |  |
| --- | --- | --- | --- | --- |
| | $W$ and $I_b$ unknown | $W$ known,<br>$I_b$ unknown | $W$ unknown,<br>$I_b$ known | $W$ and $I_b$ known |
| $n$ | locally | locally | locally | locally |
| $\text{Vol}_G$ | globally | globally | globally | globally |
| $p_1$ | nonidentifiable | globally | globally | globally |
| $p_2$ | locally | locally | locally | locally |
| $p_3$ | nonidentifiable | nonidentifiable | locally | locally |
| $p_4$ | nonidentifiable | nonidentifiable | locally | locally |
| $a_1$ | nonidentifiable | locally | nonidentifiable | locally |
| $a_2$ | locally | locally | locally | locally |
| $a_3$ | nonidentifiable | locally | nonidentifiable | locally |
| $a_4$ | locally | locally | locally | locally |
| $a_5$ | nonidentifiable | nonidentifiable | locally | locally |
| $a_6$ | globally | globally | globally | globally |
| $I_b$ | nonidentifiable | nonidentifiable | - | - |
| $W$ | nonidentifiable | - | nonidentifiable | - |

### Patient-specific parameter inference and kinetics reconstruction using OhioT1DM

In the OhioT1DM dataset, we assumed the patient’s weight is 60 kg, i.e., *W* = 60 kg. We also assume that the initial condition of the ODE system is given by 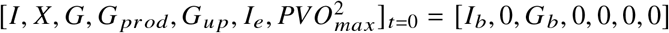, where *I_b_* is estimated from the basal insulin rate and *G_b_* is the initial blood glucose level, both of which are patient-specific estimation from the OhioT1DM dataset. We imposed the initial condition of the state variables as the auxiliary loss. We also imposed smoothing using a moving window of 30 data points on the model inputs to speed up the convergence of parameter inference. Fig. 3 shows the inference of model parameters and hidden kinetics on patient ID 588, who experienced repeated hyperglycemic events during the data collection period. Fig. 4 shows the same inference on another patient ID 591, who experienced a short period of hypoglycemia (glucose level below 80 mg/dl, we adjusted the threshold of hypoglycemia due to an overestimation of glucose levels by CGM (48)) around December 22, 2021. The corresponding results for the other four patients (IDs: 559, 563, 570, 575) can be found in Fig. S3–S6 in the Supplementary Material. Note that the time stamps in OhioT1DM dataset are pre-processed to avoid privacy leakage, hence they do not represent the real collection time. Inspired by observed fluctuation of metabolic reaction rates in oral glucose tolerance test (OGTT) (49), we adopted a time-varying parameters setting for SBINN to improve the flexibility of our model. Interestingly, while patient 588 and patient 591 followed similar insulin infusion and carb intakes, the only lifestyle difference in the exercise intensity seemingly changed the outcomes of their glucose management (Fig. S2).

**Figure 3:**
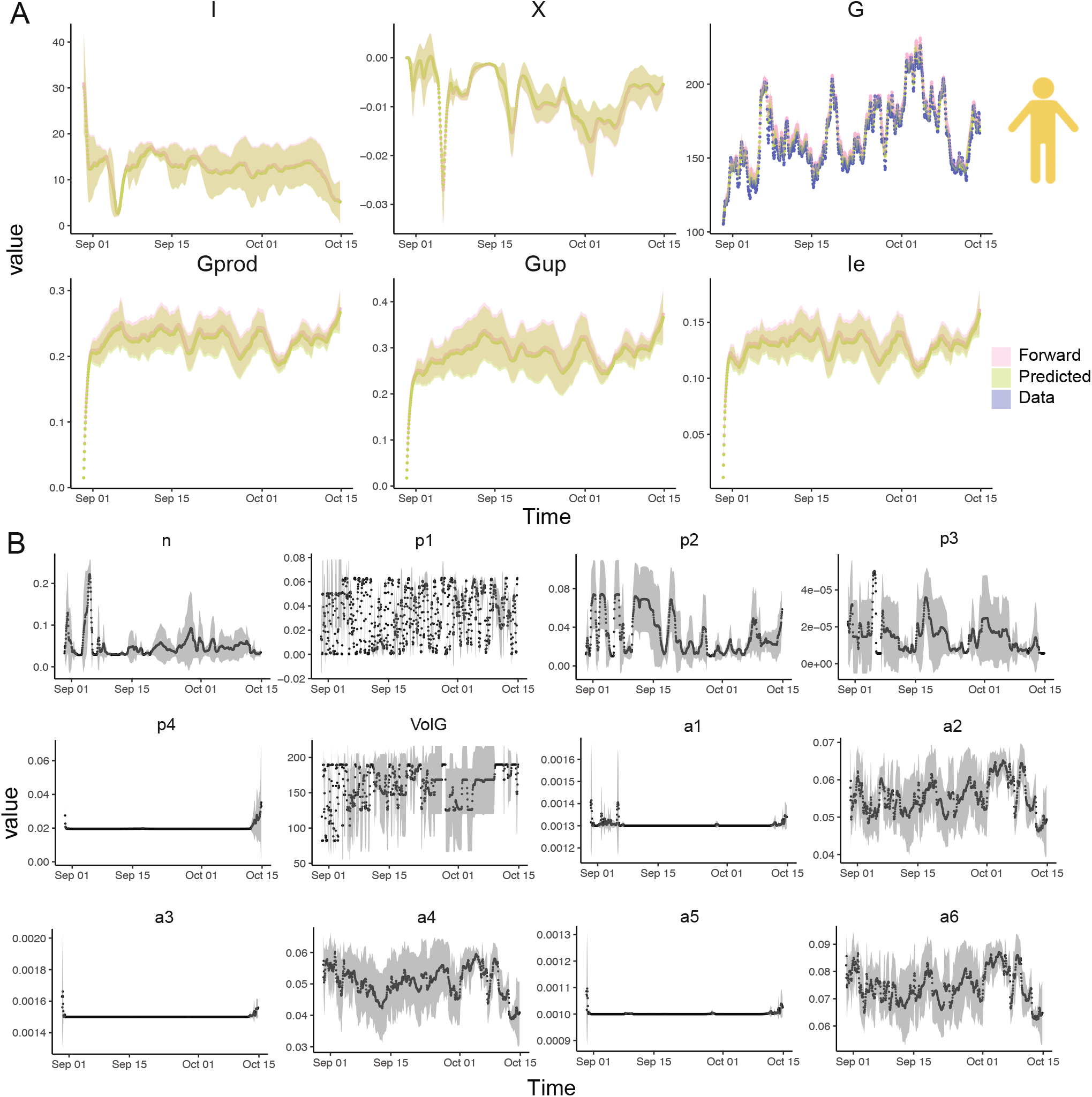
Time-dependent SBINN accurately infers the dynamics of state variables and hidden parameters in the Roy-Parker model for a patient in OhioT1DM with *recurrent hyperglycemia*. (**A**) The trajectory of state variables, plasma insulin level *I*(*t*), remote insulin level *X*(*t*), plasma glucose level *G*(*t*), exercise-induced hepatic glucose production *G_prod_* (*t*), exercise-induced glucose uptake *G_up_* (*t*), exercise-induced insulin removal from the circulatory system *I_e_* (*t*). Patient ID 588, shown here, experienced repeated hyperglycemic period, suggested by the plasma glucose levels *G* repeatedly exceeding 180 mg/dl. The blue solid dots denotes plasma glucose levels collected by CGM. The pink line denotes mean of the forward solution of the ODE from 5 different runs using inferred parameters, with the pink shade denoting the corresponding standard deviation. The green line denotes mean of denotes the predicted values by SBINN from 5 different runs using inferred parameters, with the green shade denoting the corresponding standard deviation. (**B**) The trajectory of patient-specific time-dependent hidden parameters in the Roy-Parker model for patient ID 588. The black dots denote the mean of each hidden parameters from 5 different runs and the gray shade denotes the corresponding standard deviation.

**Figure 4:**
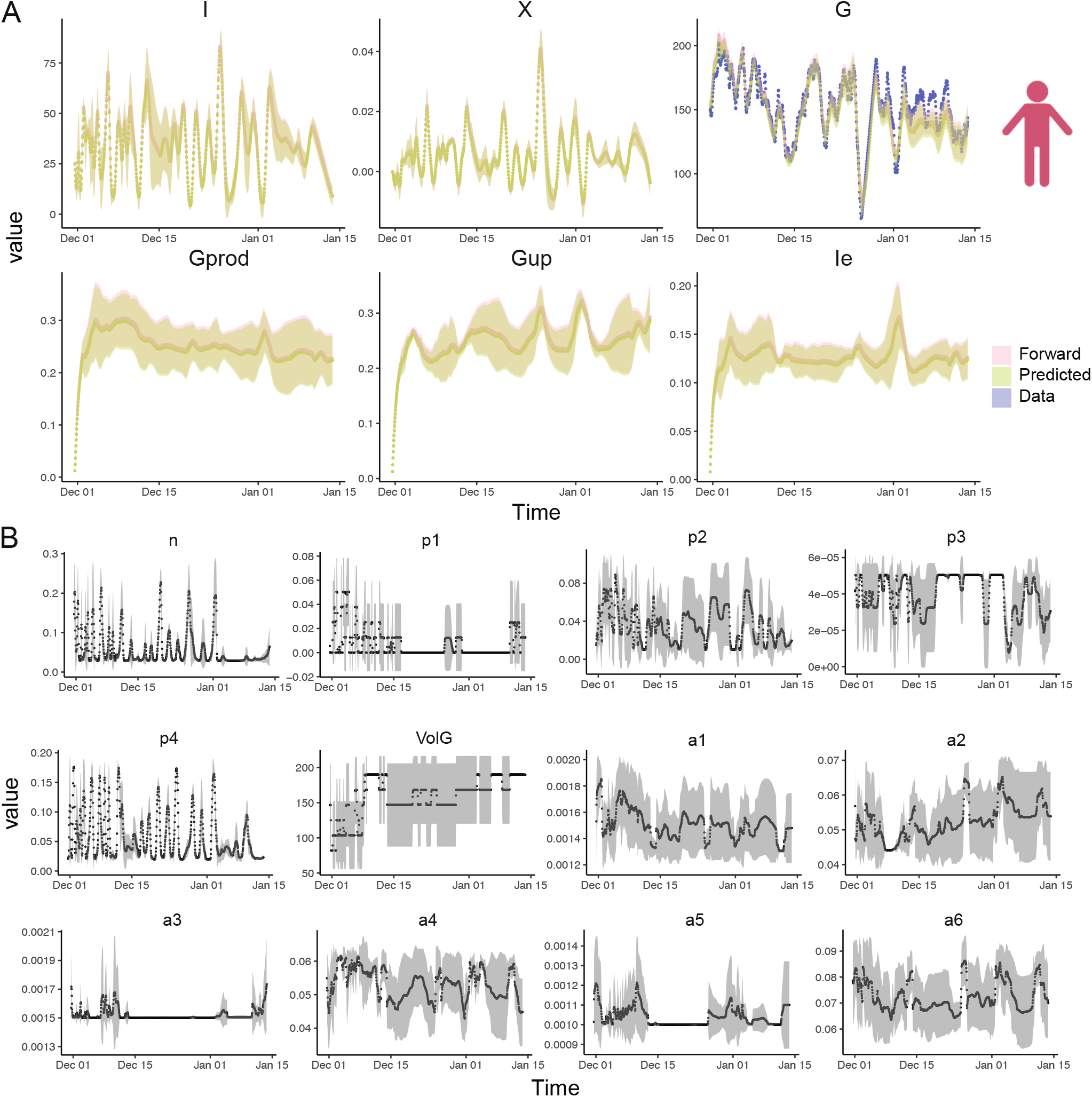
Time-dependent SBINN accurately infers the dynamics of state variables and hidden parameters in the Roy-Parker model for a patient in OhioT1DM with *occasional hypoglycemia*. (**A**) The trajectory of state variables, plasma insulin level *I*(*t*), remote insulin level *X*(*t*), plasma glucose level *G*(*t*), exercise-induced hepatic glucose production *G_prod_* (*t*), exercise-induced glucose uptake *G_up_* (*t*), exercise-induced insulin removal from the circulatory system *I_e_* (*t*). Patient ID 591, shown here, experienced a short hypoglycemic period, suggested by the plasma glucose levels went below 80 mg/dl for a short period. The blue solid dots denotes plasma glucose levels collected by CGM. The pink line denotes mean of the forward solution of the ODE from 5 different runs using inferred parameters, with the pink shade denoting the corresponding standard deviation. The green line denotes mean of denotes the predicted values by SBINN from 5 different runs using inferred parameters, with the green shade denoting the corresponding standard deviation. (**B**) The trajectory of patient-specific time-dependent hidden parameters in Roy-Parker model for patient ID 591. The black dots denote the mean of each hidden parameters from 5 different runs and the gray shade denotes the corresponding standard deviation.

We found frequently elevated plasma insulin and remote insulin in patient 591 compared to patient 588 (Fig. 3A and Fig. 4A), suggesting that patient 591 may have higher risk in developing hyperinsulinemia than patient 588. We also observed that some of the hidden parameters of patient 588 do not fluctuate as significantly over time as those of patient 591 (Fig. 3B and Fig. 4B). These parameters are *p*_4_ denoting the rate of insulin addition into the plasma from exogenous insulin, *a*_1_ denoting the rate of exercise-induced hepatic glucose production, *a*_3_ denoting the rate of exercise-induced glucose uptake, *a*_5_ denoting the rate of exercise-induced plasma insulin depletion during recovery period. Especially, we observed a sudden increase in terms of *a*_5_ for patient 591 right after the occurrence of exercise-induced hypoglycemia right before Jan 01. We also found some parameters are significantly different between patient 591 and patient 588 on average. These parameters are *n* denoting the rate of plasma insulin clearance, *p*_3_ denoting the rate of insulin addition in the remote insulin compartment and *p*_4_ denoting the rate of insulin addition into the plasma from exogenous insulin. Interestingly, all these parameters point to the balance of remote insulin and plasma insulin with *n* and *p*_3_ being almost doubled in 591 while *p*_4_ being frequently higher in 588. These findings together imply that the glucose level fluctuation is a complex process involving multiple organs and tissues, and exercise contributes to the occurrence of hypoglycemia in a more complicated way rather than merely lowering the plasma glucose level.

Our results suggest that time-depedent SBINN successfully infer the fluctuating hidden kinetics as well as the parameters for these 6 patients with high accuracy, albeit the inter-patient variability due to different daily routine of physical activities and insulin injection. More importantly, we also observed that time-dependent SBINN were able to perform robustly under 5 different random seeds with the uncertainty band confined to an acceptable range. We specifically emphasized the accuracy of our parameter inference, which helped us reconstruct the dynamics by solving a forward problem with high accuracy, indicated by the good match between pink curves (forward solver) and blue curves (real data collection).

### Offline reinforcement learning

We formulated the reinforcement learning problem for glucose regulation provided with the glucose infusion rate *u*_2_ and exercise intensity *u*_3_ from the OhioT1DM dataset, and we trained the agent to provide the optimal exogenous insulin infusion rate 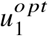 for a month. The return function depending on the resulting glucose levels at time point *t* is defined as follows,

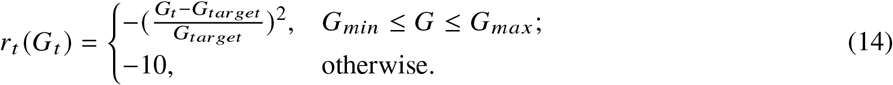

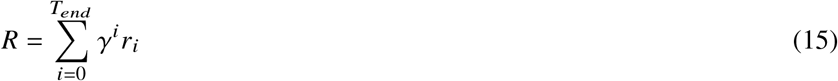

where we set the target glucose level to be 120 mg/dl, *G_min_* = 90, *G_max_* = 150. The performance of the best agent is given by the solid blue curve in Fig. 5 (A) for patient 588 and (C) for patient 591. We note here that the glucose trajectories for patient 588 and patient 591 are distinctly reflecting patients suffering from repeated hyperglycemia and hypoglycemia, respectively. Interestingly, we found that extending the permitted range of the insulin dosage for the agent by doubling the maximum action documented in the offline dataset for patient 588, who experienced repeated hyperglycemia, yielded a better glucose trajectory. When we trained the offline reinforcement learning agent, we allowed the maximum action (insulin dosage) to reach 80 *μ*U/mL, given the maximum insulin for patient 588 is around 40 *μ*U/mL. However, for patient 591 who did not have frequent hyperpglycemic events, we adopted the suggestion by Fujimoto et al. by restricting the maximum insulin dosage within the maximum insulin, i.e., 35 *μ*U/mL, recorded in the offline dataset. Figs. 5 (C) and (D) show that the offline RL agent could provide significantly better insulin dosage plans for both patient 588 and patient 591, suggested by the blue curves staying in the safe glucose region (glucose level higher than 80 mg/dl and lower than 180 mg/dl) denoted by the green shade.

**Figure 5:**
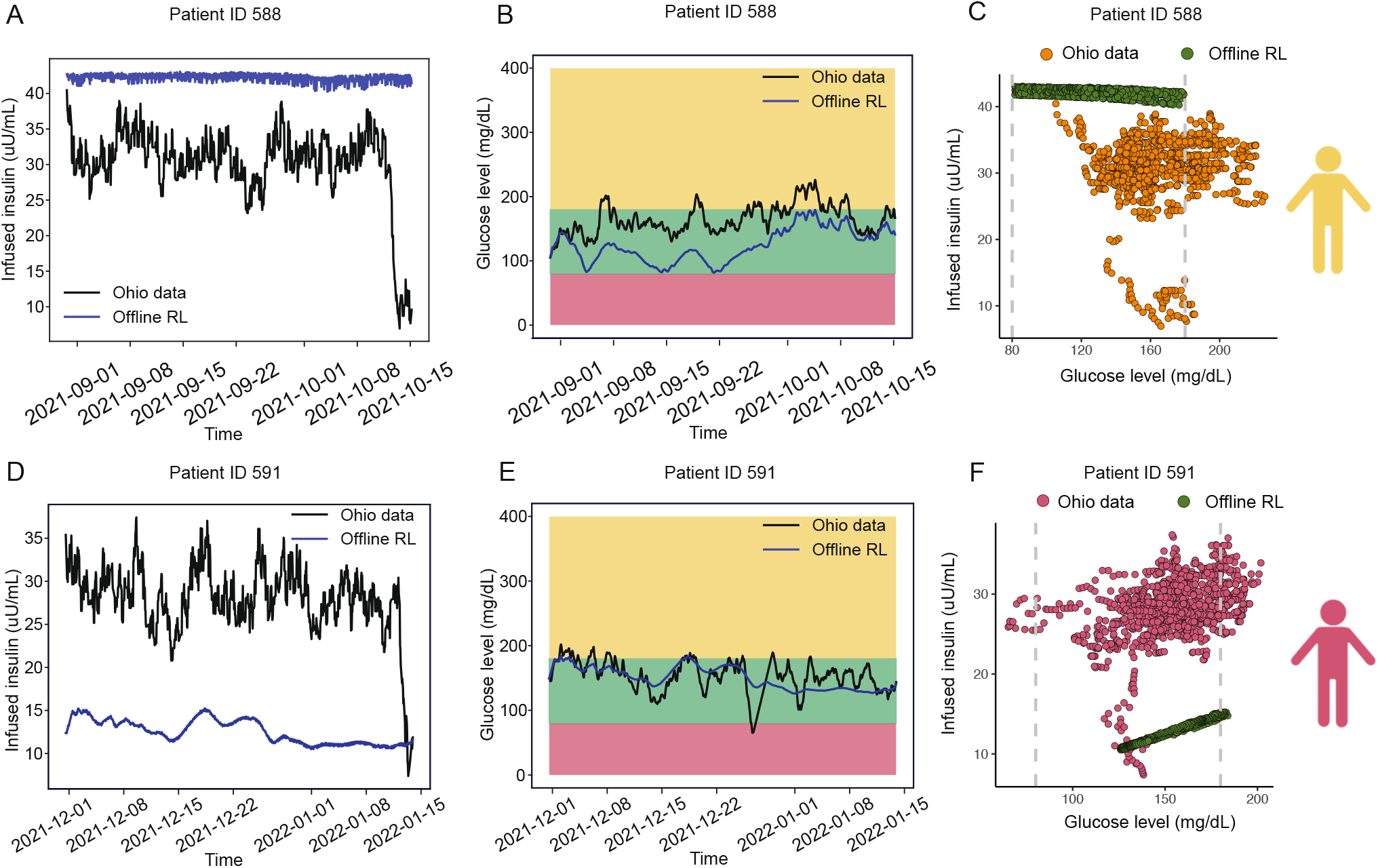
Deep offline reinforcement learning stabilizes plasma glucose levels variation and increases the time plasma glucose levels staying in normoglycemia for both patients with ID 588 and ID 591, compared to the archived data in OhioT1DM. (**A**) The solid blue curve denotes the insulin infusion rate over time provided by the offline RL agent, and the solid black curve denotes the patient’s self-operated insulin infusion rate recorded in the OhioT1DM dataset for patient ID 588. (**B**) The solid blue curve denotes the glucose trajectory using the insulin infusion plan provided by the offline RL agent for patient ID 588. The solid black curve denotes the glucose trajectory recorded in the OhioT1DM dataset for patient ID 588. Red shade denotes hypoglycemia region (BG lower than 80 mg/dl), green shade denotes normoglycemia region (BG within 80 mg/dl and 180 mg/dl), and yellow shade denotes hyperglycemia region (BG higher than 180 mg/dl). (**C**) Scatter plots of insulin infusion-glucose level pairs in OhioT1DM dataset (yellow) and the optimized offline RL (green)for patient ID 588, suggesting offline RL significantly decreases the occurrence of hyperglycemia. (**D**) The solid blue curve denotes the insulin infusion rate over time provided by the offline RL agent, and the solid black curve denotes the patient’s self-operated insulin infusion rate recorded in the OhioT1DM dataset for patient ID 591. (**E**) The solid blue curve denotes the glucose trajectory using the insulin infusion plan provided by the offline RL agent for patient ID 591. The solid black curve denotes the glucose trajectory recorded in the OhioT1DM dataset for patient ID 591. Red shade denotes hypoglycemia region (BG lower than 80 mg/dl), green shade denotes normaglycemia region (BG within 80 mg/dl and 180 mg/dl), and yellow shade denotes hyperglycemia region (BG higher than 180 mg/dl). (**F**) Scatter plots of insulin infusion-glucose level pairs in OhioT1DM dataset (yellow) and the optimized offline RL (green) for patient ID 588, suggesting offline RL prevents the dangerous hypoglycemia and decreases the time of hyperglycemia.

## 4 CONCLUSION

Insulin is the mainstay of treatment for patients with type 1 diabetes mellitus and long-standing type 2 diabetes mellitus to achieve good glycemic control (50). Overestimation of the necessary insulin dosage can be extremely dangerous and may lead to fatally low blood glucose levels below 80 mg/dl when measured by CGM, namely hypoglycemia, while underestimated insulin leaving blood glucose above 180 mg/dl may result in hyperglycemia, which is believed to be responsible for micro- and macrovasular diseases in the long run. In modern medicine, the use of insulin pumps along with continuous glucose monitors has made it easier but requires significant resources and patient education. Fortunately, a closed-loop control system, also called artificial pancreas, which automates insulin infusion so as to maintain a consistently stable blood glucose level, undoubtedly relieves the burden of both patients and doctors and saves medical costs.

In this work, we attempted to address a few of the challenges with our framework, which effectively combines three key components to build a patient-specific artificial pancreas simultaneously considering data collected from wearable devices, i.e., meal intake, insulin infusion and physical exercises. These important components are: 1) a real-world historical medical dataset, namely OhioT1DM dataset, containing patient-specific glucose, insulin, meal intake and exercise intensity; 2) a flexible ODE model defining glucose-insulin dynamics by systematically prioritizing two significant external factors, i.e., meal intake and physical exercise intensities; and 3) an offline reinforcement learning algorithm, namely batch constrained Q-learning, without directly interacting with the patient’s metabolic environment. With time-dependent systems biology informed neural networks (SBINN) along with patient-specific data, our model not only correctly predicts the hidden states that cannot be measured with current diabetes technology, but also accurately infers patient-specific parameters governing the patient-specific Roy-Parker model by reconstructing all states of interest. More importantly, the uncertainty quantified hidden parameters provide patient-specific clinical interpretations on how patients’ behavioral pattern shape their corresponding glucose-insulin dynamics and can contribute to the prognostic suggestion on future patient-specific behaviors. Furthermore, we were able to train an agent to automate insulin infusion and optimize the performance of the agent on the patient-specific ODE model with same glucose infusion and exercise intensity over time. Our results suggest that the best trained agent has a better performance in terms of maintaining blood glucose levels within the safe range, compared to the self-operated insulin infusion by patients themselves. This suggests that an agent trained by offline reinforcement learning is able to learn a better insulin dosage depending solely on past glucose level sequences without meal or exercise announcements, implying the advantageous potential of offline reinforcement learning in translational healthcare.

In spite of the improved glycemic control provided by our offline agent and minimal human intervention demanded by our framework, we can still identify possible improvements to the proposed framework, considering ODE model development, disease characterization and data processing. As is believed that there are both a time delay in the effect of insulin on the glucose production and that on the glucose utilization (27), it may be helpful to modify the ODE model to address the sluggish effects. Additionally, given that insulins are categorized into fast-acting, intermediate-acting and long-acting depending on the timing of their action in body, we could extend the data preprocessing of external insulin as well as update the ODE model to allow variations in insulin types (51, 52). Another avenue to be considered in improving closed-loop systems is by adding amylin or glucagon, a hormone also produced by the pancreas, resulting to a multi-hormone closed-loop system (53). With glucagon being the additional action option, the reinforcement learning agent will be able to explore insulin action space with a higher degree of freedom.

Despite of the aforementioned possible improvements, our framework shows inspiring translational potentials in precision medicine by enriching the limited options of digital models utilizing medical data collected from wearable devices. For example, with a few slight modifications on the ODE model to address the insulin resistance of tissue, we could model and optimize the insulin usage in patients with type 2 diabetes who develop insulin resistance (54). Interestingly, novel wearable devices are being designed to probe insulin and other metabolites that are impossible to quantify with existing techniques (55, 56), which will clearly benefit the calibration and the extension of our framework. Furthermore, the rapid development in non-invasive wearable devices tremendously increases patients’ compliance to the use of wearable devices (57, 58), and hence provides data with more frequent readouts, higher resolution and diverse modalities, which will obviously enhance the performance of our framework beyond diabetes related research. In summary, by seamlessly integrating real-world data from wearable sensors, systems biology informed numerical models and offline reinforcement learning algorithms, we believe our novel framework could shed light on the translational precision medicine where adequate biological numerical models are established and enough amount of medical data from wearable devices are available.

## 5 DATA AVAILABILITY

The OhioT1DM dataset used in the current study is publicly available on http://smarthealth.cs.ohio.edu/OhioT1DM-dataset.html.

## 6 CODE AVAILABILITY

The code will be released upon acceptance of publication.

## ACKNOWLEDGEMENTS

G.E.K. would like to acknowledge support by NIH U01HL142518 and R01HL154150. C.S.M. would like to acknowledge support by NIH DK081913.

## Author Contributions

G.E.K. and C.S.M. supervised the work and formulated the problem. Y.D. and G.E.K. developed the model. Y.D. implemented the computer code. Y.D. performed computations. Y.D. and G.E.K. analyzed data. Y.D., K.A., C.S.M., and G.E.K. wrote the paper.

## 7 Supplementary Material

**Table S1:**
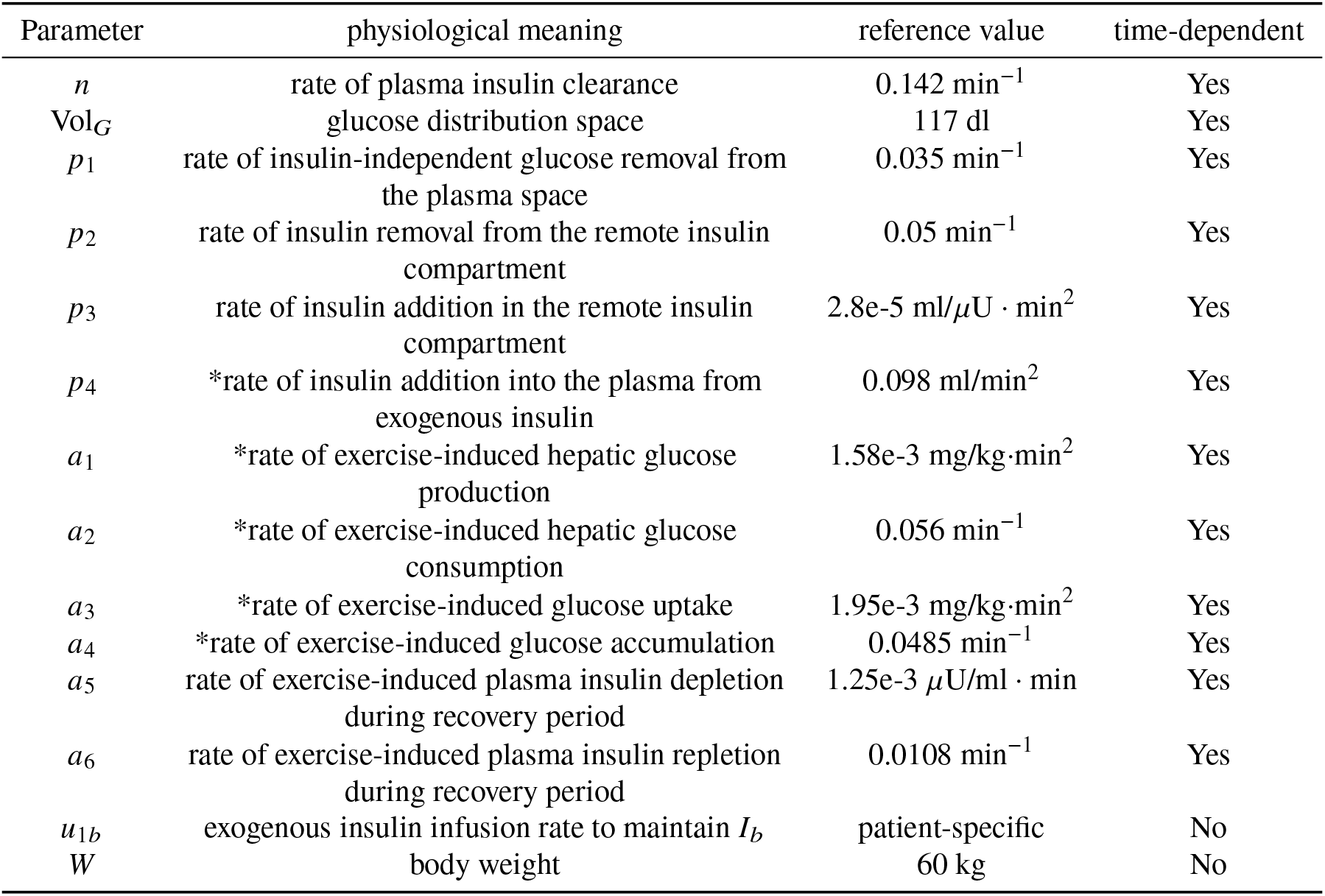
Nomenclature and physiological meanings of patient-specific parameters in the Roy-Parker model. Physiological meanings marked with * were not explained in the original publication, hence standing for the interpretation given the ODE form.

### Patient record example

**Figure S1:**
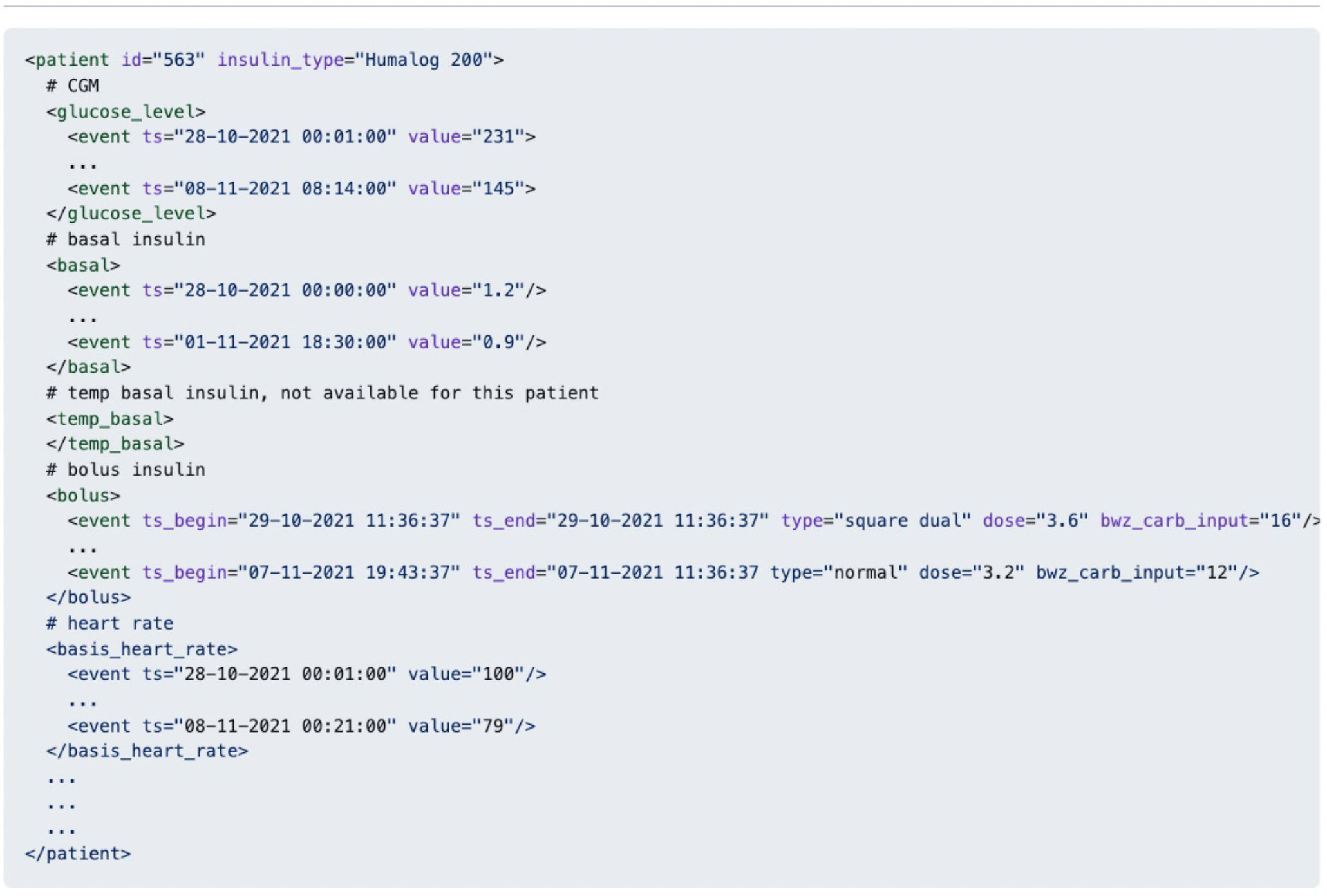
A pseudo example of patient’s record in the OhioT1DM dataset, showing the structure of the tabular data.

**Figure S2:**
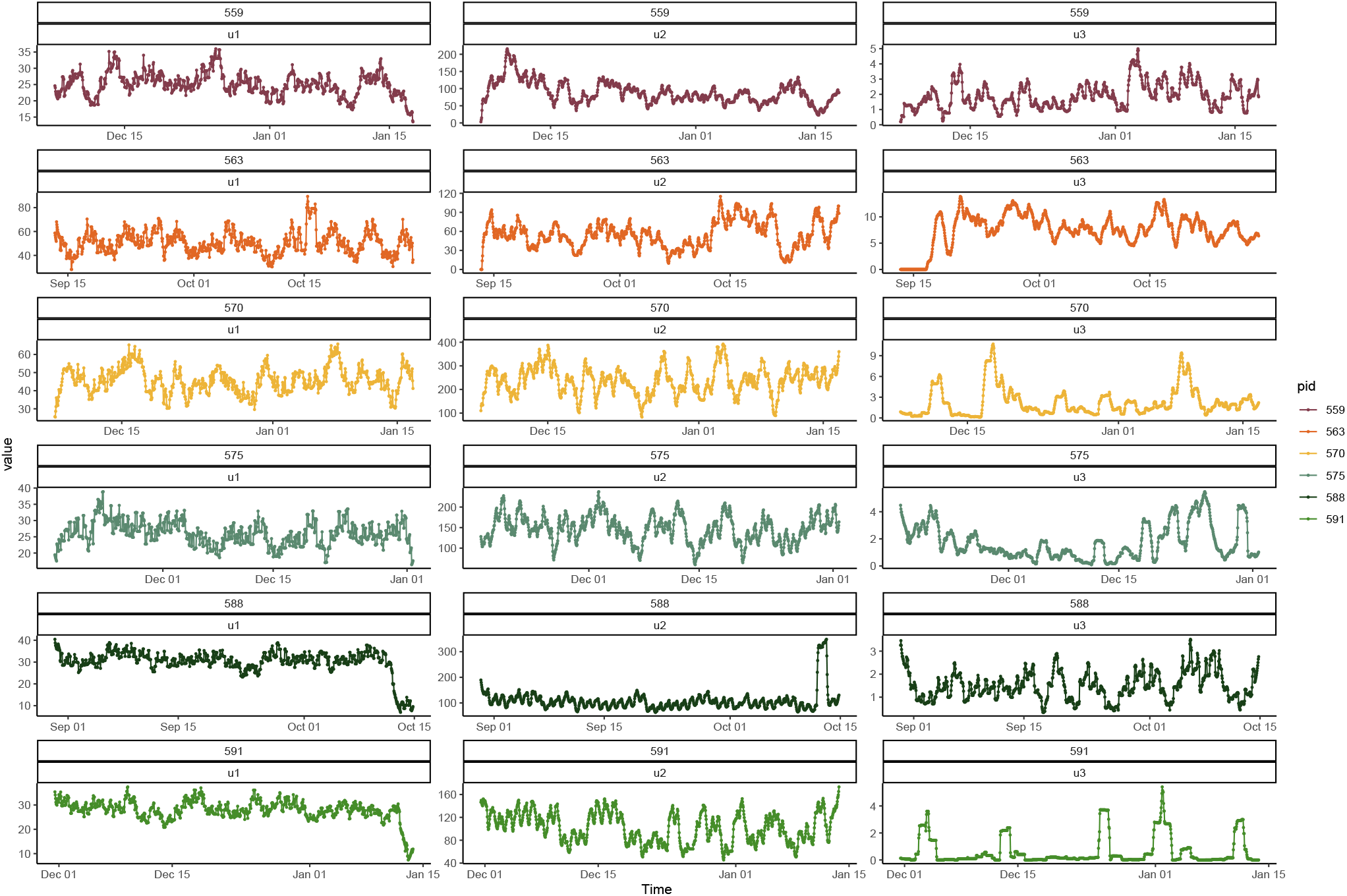
External inputs extracted from patients’ record in the OhioT1DM dataset. *u*_1_ denotes insulin infusion; *u*_2_ denotes carbohydrate intakes; *u*_3_ denotes exercise intensity.

**Algorithm 1.**
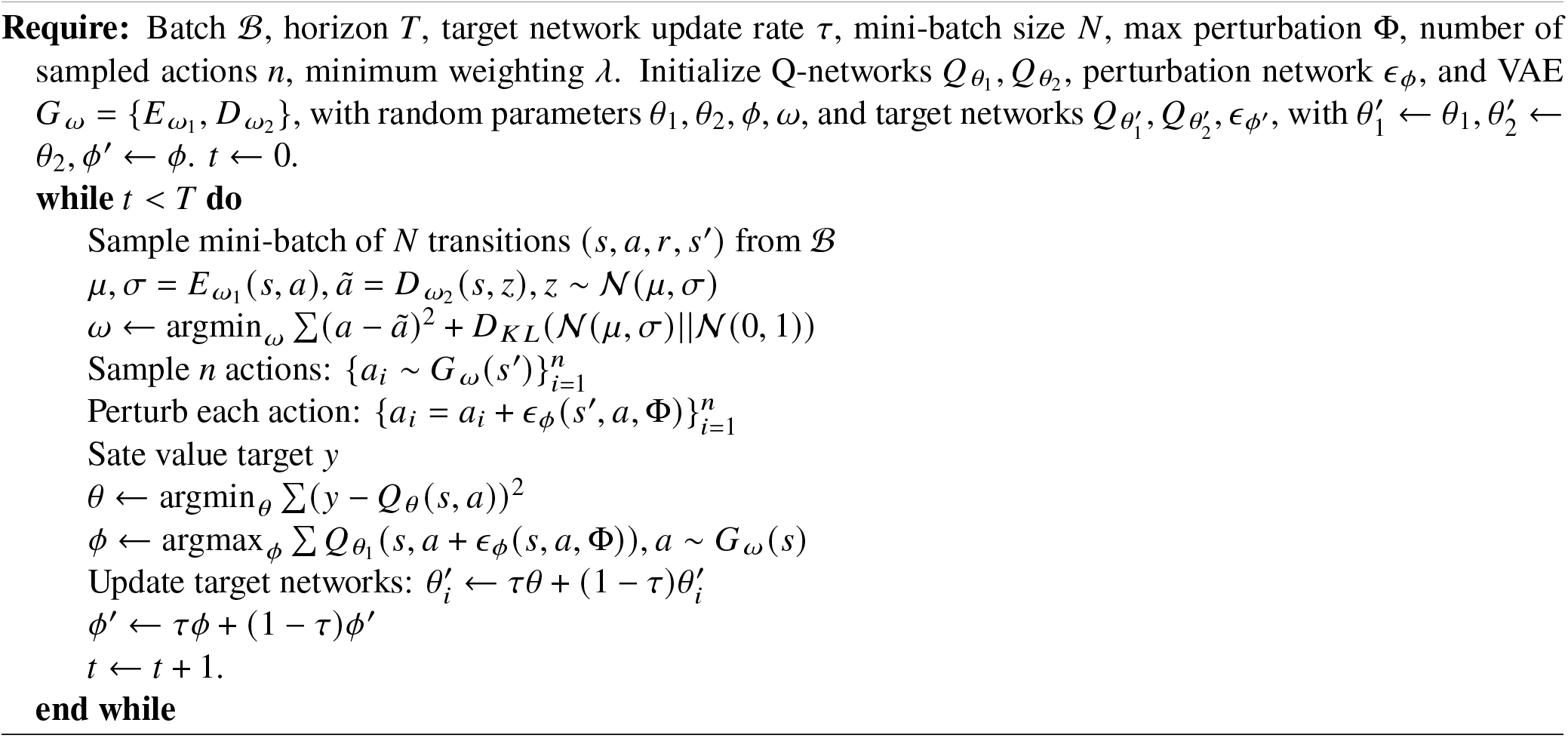
Batch constrained Q-learning algorithm.

**Table S2:**
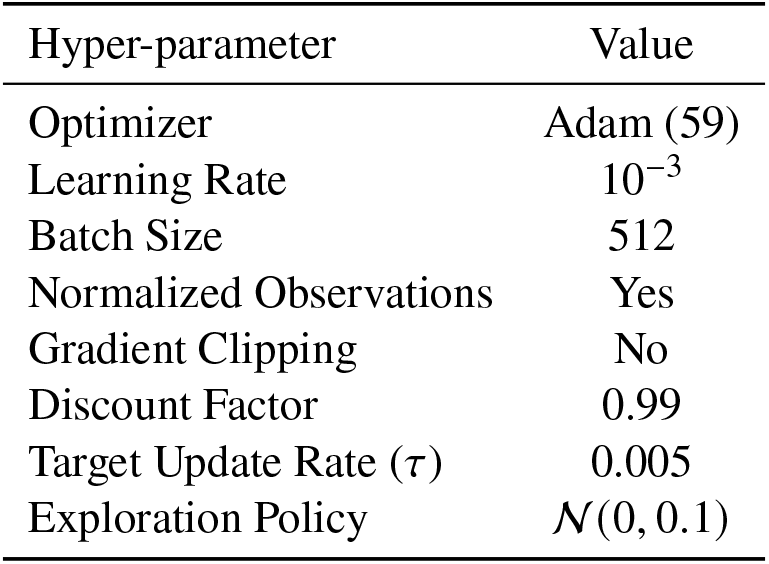
Hyper-parameters used in the deep offline reinforcement learning.

**Figure S3:**
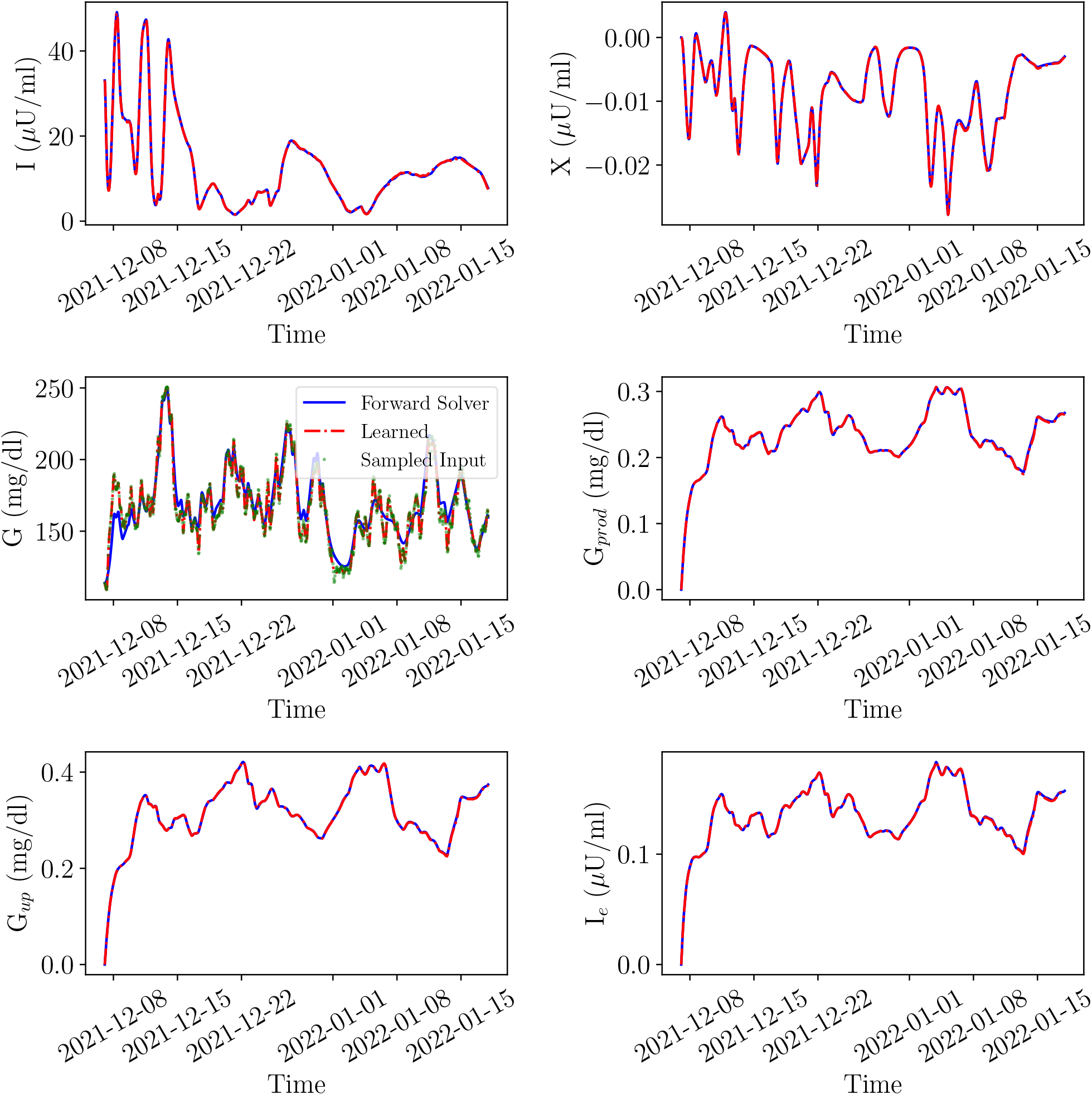
Prediction of 6 state variables in the Roy-Parker model using SBINNs patient with ID 559.

**Figure S4:**
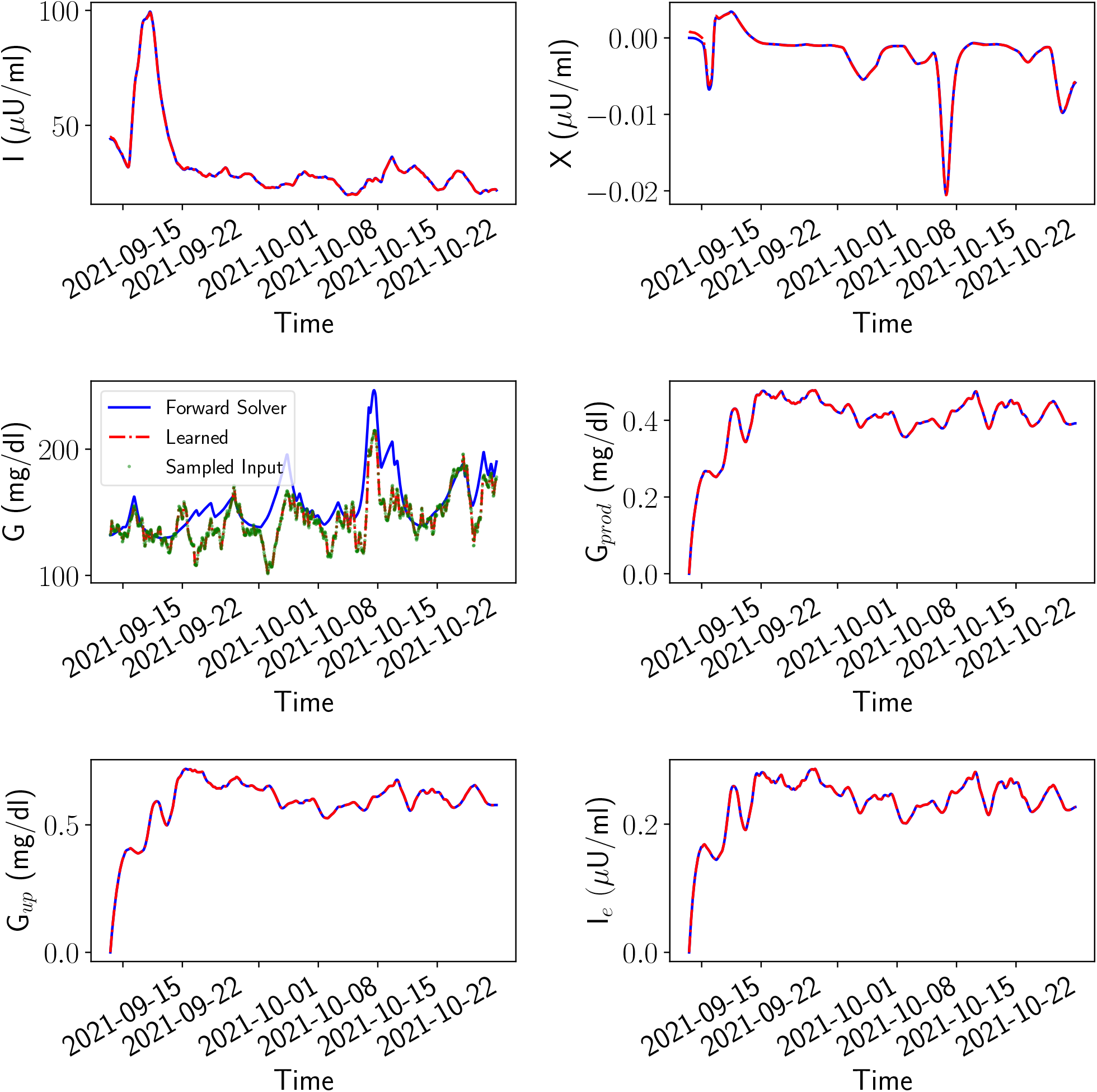
Prediction of 6 state variables in the Roy-Parker model using SBINNs patient with ID 563.

**Figure S5:**
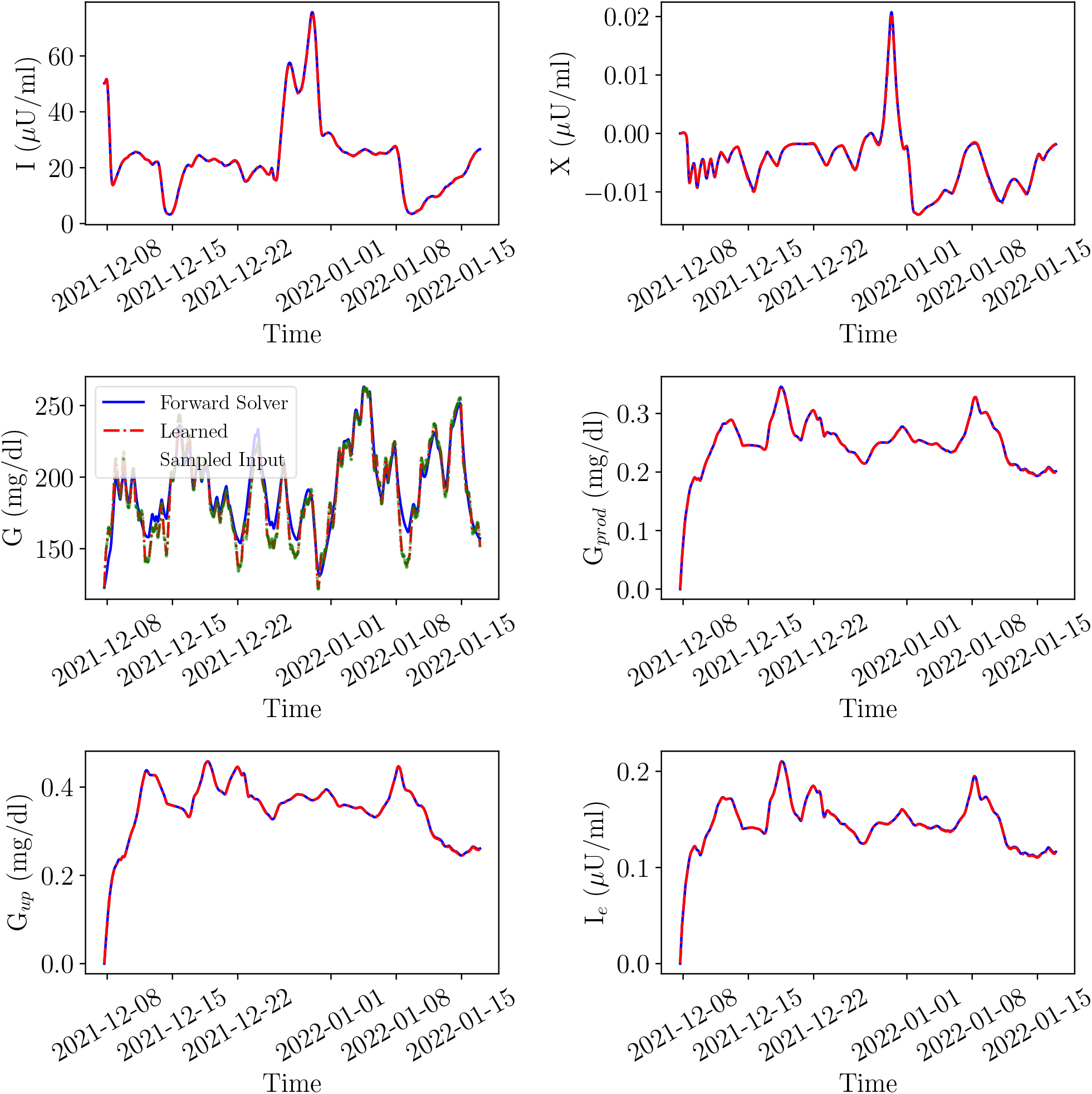
Prediction of 6 state variables in the Roy-Parker model using SBINNs patient with ID 570.

**Figure S6:**
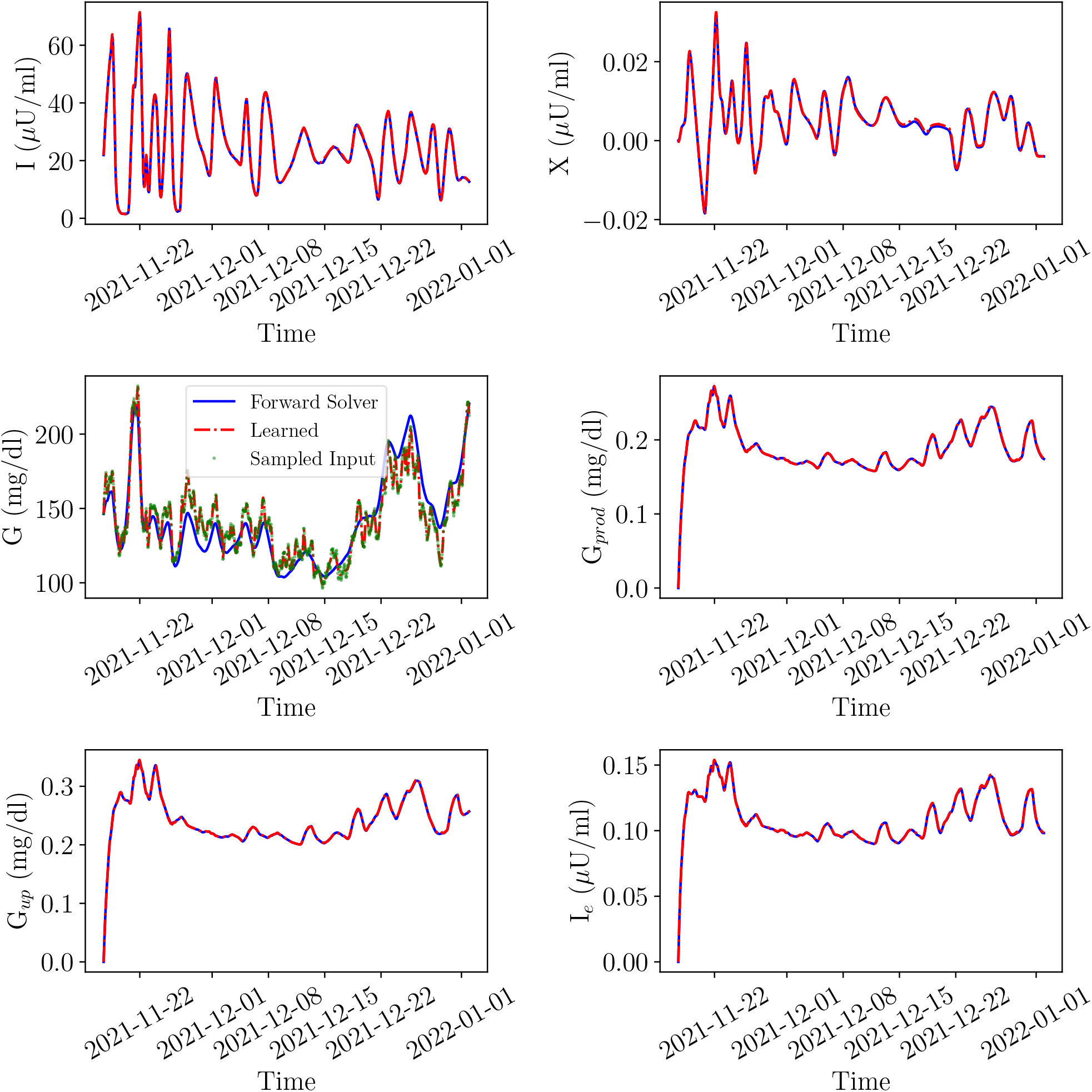
Prediction of 6 state variables in the Roy-Parker model using SBINNs patient with ID 575.

